# Targeted metagenomic recovery of coronaviruses from wildlife samples

**DOI:** 10.64898/2026.05.05.722935

**Authors:** Camille Melissa Johnston, Vithiagaran Gunalan, Hans J. Baagøe, Morten Elmeros, Esben T. Fjederholt, Michelle Lauge Quaade, Mie J. S. Hansen, Louise Lohse, Thomas Bruun Rasmussen

## Abstract

Coronaviruses (CoVs) are diverse RNA viruses infecting a wide range of hosts, with significant implications for public health, animal production and welfare. Bats are key reservoirs of mammalian CoVs and have contributed to the emergence and circulation of several zoonotic viruses in a wildlife context, while mustelids represent important hosts at the human-animal interface, as highlighted by SARS-CoV-2 outbreaks in farmed mink.

Using metagenomic next-generation sequencing, we screened bats and wild mustelids and recovered and characterized full-length CoV genomes. Building on previous findings in Danish wildlife, including alpha-CoVs and a MERS-related beta-CoV in bats, this study expands current knowledge of coronavirus diversity and evolution in wildlife, emphasizing their relevance for zoonotic emergence.

## Introduction

Coronaviruses (CoVs) are RNA viruses associated with a wide range of respiratory and enteric infections in humans and animals. Their ability to infect diverse host species underlines their importance from a One Health perspective, due to their impact on public health and livestock-associated disease with substantial economic consequences (Corman et al., 2018; V’kovski et al., 2021). Members of the family *Coronaviridae* (order *Nidovirales*) are classified into four genera: Alpha-, beta-, gamma-, and delta-CoVs. Alpha- and beta-CoVs predominantly infecting mammals, whereas gamma- and delta-CoVs circulate widely in avian hosts (Woo et al., 2023).

Bats (order Chiroptera) are considered key reservoirs of mammalian coronaviruses, as they harbor a remarkably high genetic diversity of both alpha- and beta-CoVs. Phylogenetic evidence indicates that bats have played a central role in the evolutionary history of several human and animal CoVs, including the progenitors of severe acute respiratory syndrome (SARS) CoV and SARS-CoV-2, and possibly Middle East respiratory syndrome (MERS) CoV (Anthony et al., 2017; Hu et al., 2015; Memish et al., 2013; Temmam et al., 2022). Bat-associated alphacoronaviruses have also been implicated in the emergence of human CoVs such as HCoV-229E and HCoV-NL63, as well as several livestock pathogens, e.g. porcine epidemic diarrhea virus and transmissible gastroenteritis virus (Corman et al., 2018; Huang et al., 2013; Yang et al., 2020).

In addition to bats, mustelids (family Mustelidae) represent an important group of CoV hosts, particularly at the human-animal interface. Experimental infection have demonstrated the susceptibility of ferrets (*Mustela putorius furo*) to SARS-CoV and SARS-CoV-2, and large-scale outbreaks of SARS-CoV-2 have been documented in farmed American mink (*Neogale vison*) following transmission from humans, with evidence of subsequent spill-back events (Boklund et al., 2021; Cahan, 2020; Kim et al., 2020; Larsen et al., 2021; Martina et al., 2003; Molenaar et al., 2020; Oreshkova et al., 2020; Oude Munnink et al., 2021; Rasmussen et al., 2024; Richard et al., 2020; Schlottau et al., 2020). Several alpha-CoVs belonging to the subgenus *Minacovirus*, have been described in ferrets, American minks, and stoats (*Mustela erminea*) (Apaa et al., 2023; Lazov et al., 2025; Martínez et al., 2008; Quaade et al., 2026; Vlasova et al., 2011; Williams et al., 2000) and additional CoVs have recently been detected in other mustelid species, such as a possibly new genus epsilon-CoV in Italian badgers (*Meles meles*) and possible gamma-CoV in Chinese ferret badgers (*Melogale moschata*) (Dong et al., 2007; Zamperin et al., 2023).

Coronaviruses are enveloped viruses with large, linear, positive-sense single-stranded RNA genomes of approximately 27–31 kb. The first two-thirds of the genome encodes the viral replicase genes through two partially overlapping open reading frames (ORFs), ORF1a and ORF1b. Translation of ORF1b depends on a ribosomal -1 frameshift, resulting in the synthesis of two polyproteins of different lengths. These polyproteins are subsequently cleaved by virus-encoded proteases into 15-16 non-structural proteins. The last third of the genome encodes the structural and accessory proteins. All coronaviruses possess four principal structural proteins: spike (S), envelope (E), membrane (M), and nucleocapsid (N), which are typically arranged in the order S-E-M-N. Additional accessory genes are variably interspersed among the structural genes and at the 3’ end of the genome (de Groot et al., 2012; Lu et al., 2020; Sola et al., 2015; V’kovski et al., 2021).

Metagenomic next-generation sequencing (mNGS) has become an important tool for virus discovery and genome characterization, enabling comprehensive detection of viral sequences present in biological samples (Nooij et al., 2018). In addition to facilitating recovery of complete CoV genomes, mNGS can reveal the broader viral diversity present in wildlife samples, as demonstrated by the recovery of full-length astrovirus genomes from bats (Lazov et al., 2021).

Using this mNGS approach, we have previously recovered full-length alpha-CoV genomes from multiple Danish bat species and farmed American mink, and reported the first detection and complete genome of a MERS-related beta-CoV in brown long-eared bats (*Plecotus auritus*) (Johnston et al., 2025; Lazov et al., 2021, 2025). Here, we extend this work by screening bats and wild mustelids for CoVs and applying an optimized metagenomic workflow to recover and characterize full-length viral genomes, thereby expanding our understanding of coronavirus diversity in wildlife.

## Materials and Methods

### Collection of samples

Dead-found mustelids were collected as part of a surveillance project aimed at elucidating the health status and occurrence of selected pathogens in native mustelid species in Denmark (Elmeros et al., 2025). Most specimens were collected between 2022 and 2024 and primarily consisted of road-killed animals, while a smaller number were collected in between 2019 and 2021 (**Figure 1** and **Table S1**). Fecal and throat swabs were collected from each specimen.

**Figure 1.**
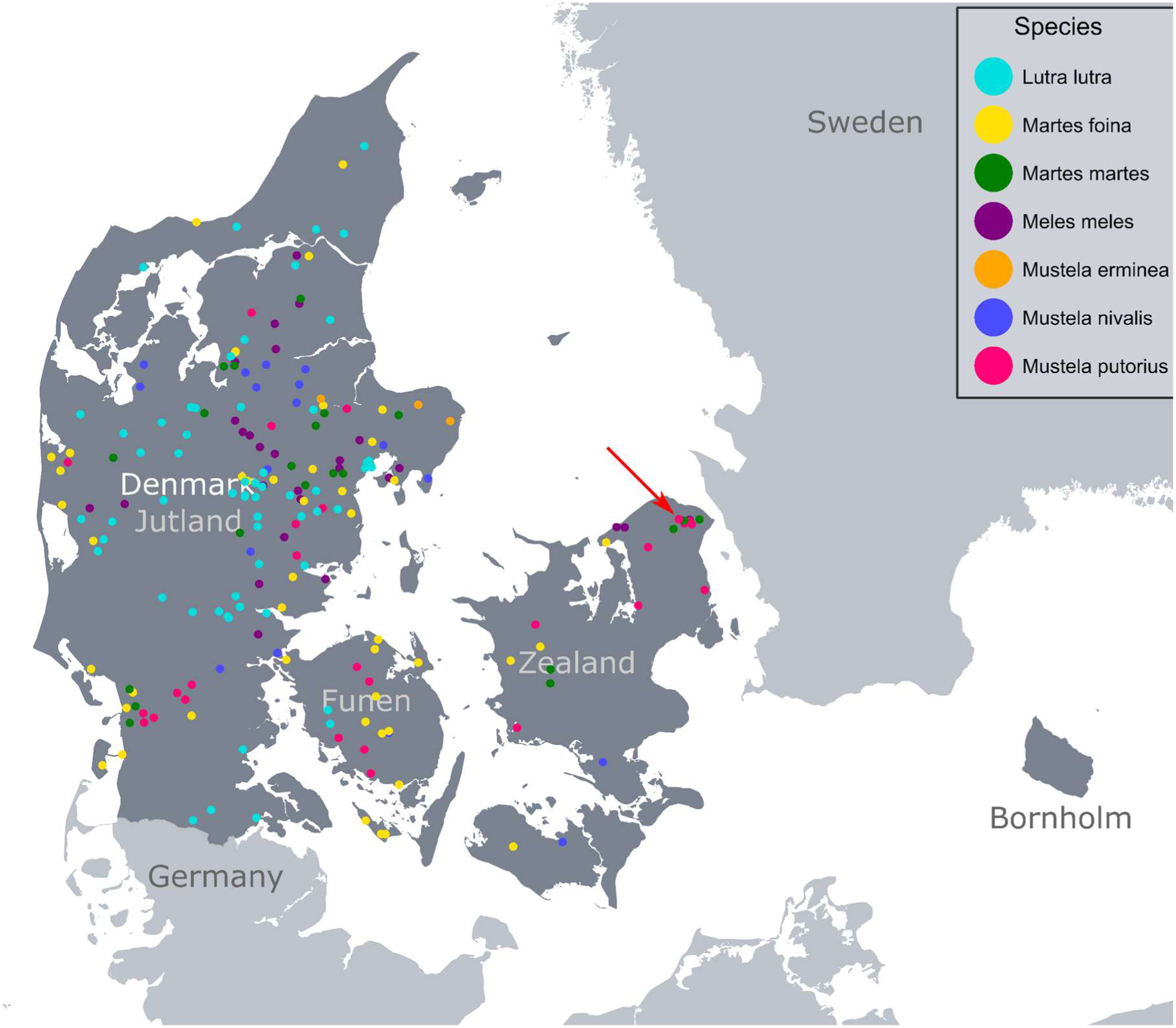
Geographic distribution of sampled mustelids across Denmark. *Lutra lutra* (cyan), *Martes foina* (yellow), *Martes martes* (green), *Meles meles* (purple), *Mustela erminea* (orange), *Mustela nivalis* (cobalt blue), and *Mustela putorius* (hot pink). The red arrow indicates the location of the single pan-CoV positive mustelid sample, detected in *M. putorius*.

Fecal samples from live bats and fecal or colon samples from whole carcasses from thirteen different bat species were collected as part of a national surveillance program of bats in Denmark for the years 2021-2024. The program included both active surveillance, consisting of non-invasive collection of fecal samples from live bats, and passive surveillance, based on sampling of dead individuals submitted from multiple geographic regions (**Figure 2** and **Table S2**). Sampling from live bats was conducted with minimal disturbance to the animals, and associated metadata such as species identification, location, and date of collection were systematically recorded.

**Figure 2.**
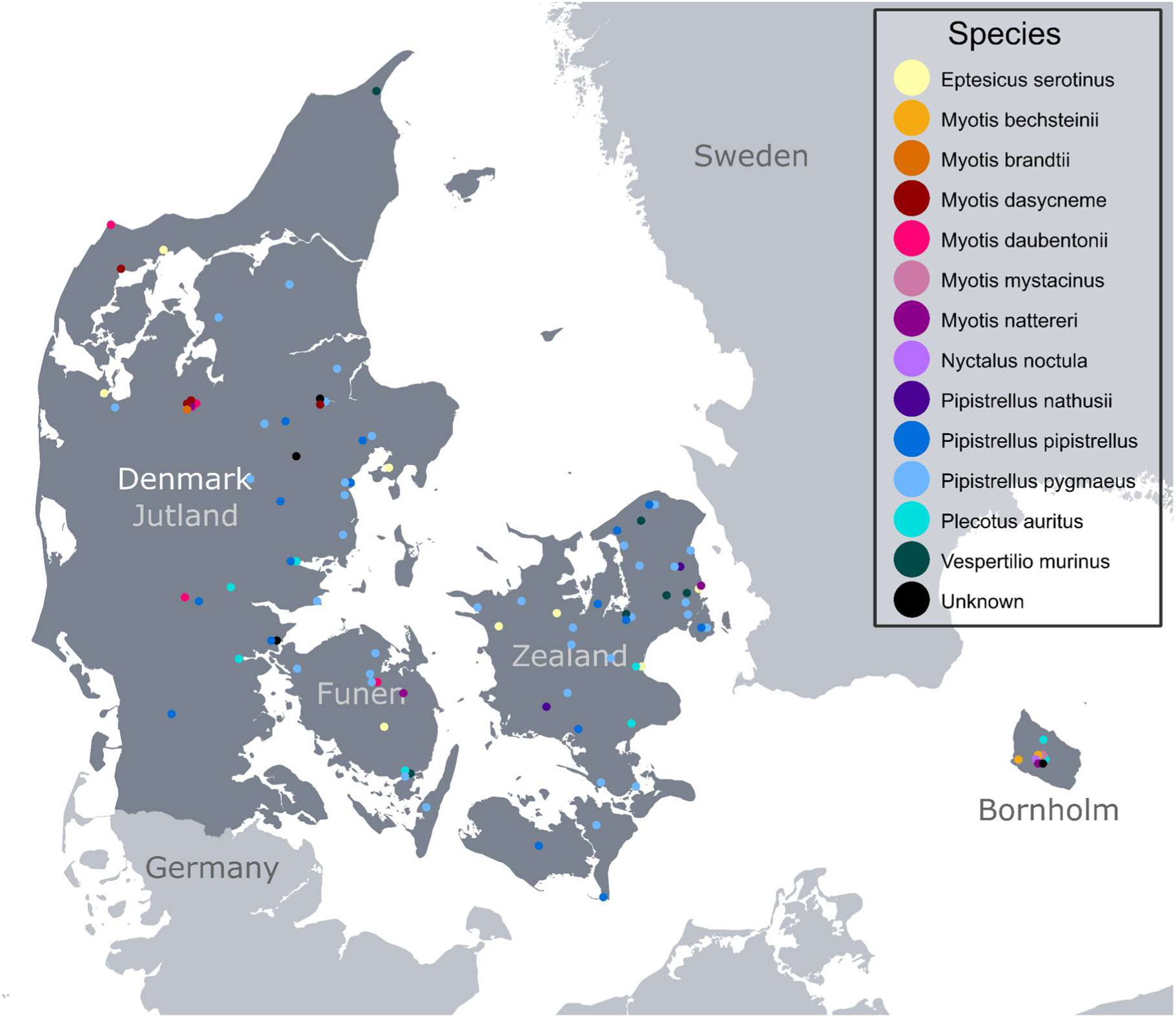
Geographic distribution of bat sampling locations across Denmark. Points represent unique species-location combinations after filtering for duplicate records. Where multiple species were recorded at the same location, jitter was applied for visualization purposes. *Eptesicus serotinus* (pale yellow), *Myotis bechsteinii* (golden yellow), *Myotis brandtii* (burnt orange), *Myotis dasycneme* (dark red), *Myotis daubentonii* (hot pink), *Myotis mystacinus* (dusty rose), *Myotis nattereri* (dark magenta), *Nyctalus noctula* (lavender), *Pipistrellus nathusii* (deep violet), *Pipistrellus pipistrellus* (cobalt blue), *Pipistrellus pygmaeus* (light blue), *Plecotus auritus* (cyan), *Vespertilio murinus* (dark teal), and unknown species (black).

### Host species determination

Live bats were identified to species level based on morphological characters. Bats from passive surveillance were identified by PCR amplification and sequencing of mitochondrial 16S RNA and cytochrome b genes (**Table 1**), as previously described (Lazov et al., 2018).

**Table 1:**
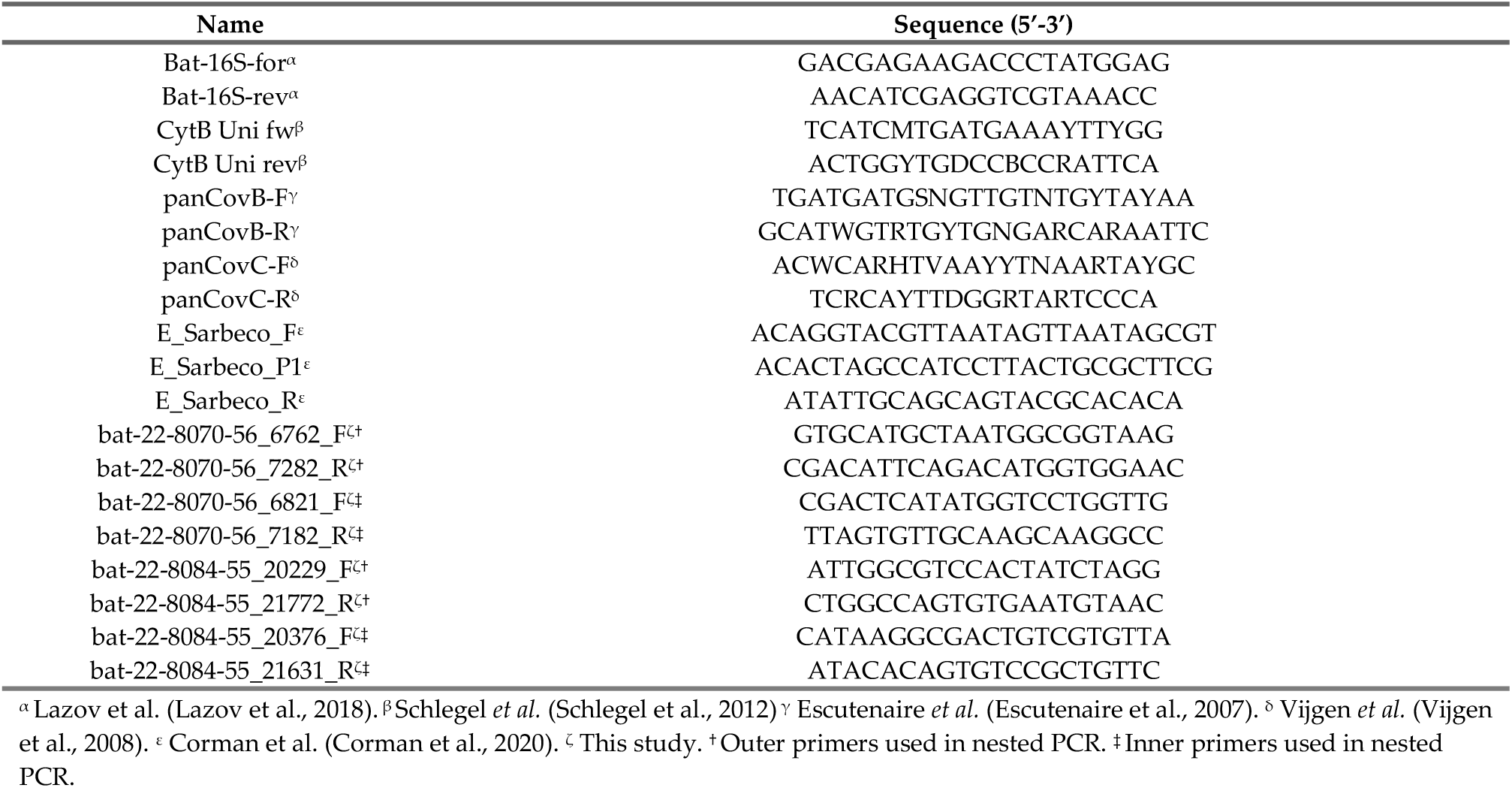
Primer overview.

### Screening of samples with pan-CoV RT-PCR assays

Extracted RNA from fecal and colon samples from bats and mustelids were screened using two pan-CoV RT-PCR assays, panCoV B and panCoV C (**Table 1**), and selected pan-CoV positive samples were subjected to Sanger sequencing of the corresponding amplicons, as previously described (Johnston et al., 2025; Lazov et al., 2018).

### Screening of samples with SARS-CoV-2 RT-qPCR assay

Fecal and colon samples from bats and throat samples from mustelids were screened using SARS-CoV-2 RT-qPCR assay, as previously described (Corman et al., 2020; Hammer et al., 2021), using a modified extraction protocol. Briefly, viral RNA was extracted from the fecal, colon and throat samples using a MagNA Pure 96 system (Roche, Basel, Switzerland) with the DNA and Viral NA Small Volume Kit and the Viral NA Plasma external lysis S.V. 3.1. protocol. Prior to extraction the fecal and colon samples were homogenized in Eagle’s medium using a TissueLyser II (QIAGEN, Hilden, Germany) and centrifuged. Extracted RNA samples were screened for the presence of SARS-CoV-2 using primers targeting the E gene (**Table 1**). The products were amplified by RT-qPCR using Luna Probe One-Step RT-qPCR with the E gene primers in a final volume of 20 µL, containing 0.4 µM of each primer and 0.2 µM of the probe. The PCR program was as follows: 55°C for 10 min, followed by 95°C for 3 min, then 45 cycles of 95°C for 15 s and 58°C for 30 s.

### Metagenomic workflow

Samples positive in the pan-CoV assays were subjected to sequencing. Double-stranded cDNA was generated as previously described (Johnston et al., 2025; Lazov et al., 2021). The resulting cDNA was sequenced on a MiSeq platform (Illumina, San Diego, CA, USA) with a modified Nextera XT DNA library protocol and the MiSeq reagent kit v2 (300 cycles), generating 2x150 bp paired-end reads. Selected samples were deep sequenced on a NextSeq 1000 system (Illumina) using a modified DNA Prep library protocol with the XLEAP-SBS 300 cycles reagent kit, producing 2x150 bp paired-end reads. For samples sequenced on both platforms, reads were combined prior to downstream analysis.

Reads were trimmed using Trim Galore (Krueger et al., 2023), a wrapper for Cutadapt (Martin, 2011), for quality (q26). Trimmed reads were taxonomically classified using both Kraken2 (Wood et al., 2019) with the plusPFP database and DIAMOND (Buchfink et al., 2015) against the RefSeq protein database (Buchfink et al., 2021; Wood et al., 2019). Trimmed reads classified as viral or unknown were retained for *de novo* assembly using both MEGAHIT v.1.2.9 (D. Li et al., 2015) and metaSPAdes v4.0.0 (Nurk et al., 2017). Resulting contigs were filtered for a minimum length of 500 bp and queried against a local BLAST *nt_viruses* database using BLASTn (Camacho et al., 2009). For each contig, the best reference hit was selected based on the lowest E-value, highest bitscore and highest percent identity. Contigs were then mapped using minimap2 (H. Li, 2018) against a combined FASTA file comprising all best reference hits. Coverage statistics and visualizations were generated in R using Qualimap v2.3 (García-Alcalde et al., 2012), bamsignals v1.34.0 (Mammana & Helmuth, 2023), ggplot2 v3.5.1 (Wickham, 2016), ggplotify v0.1.2 (Yu, 2023) and plotly v4.10.4 (Sievert, 2020).

For draft genome assembly, contigs were grouped based on taxonomic assignment at the family level (e.g. *Coronaviridae*). The most frequently occurring best reference hit among the contigs in each group was selected as a scaffold. All contigs were mapped to this reference using minimap2, and a consensus sequence was extracted with Samtools (Danecek et al., 2021). This consensus was polished with trimmed reads using BWA-MEM v0.7.18 (H. Li, 2013). In addition, a purely reference-based consensus sequence was generated by mapping trimmed reads directly to the reference. Gap closing was performed using two strategies in parallel: I) extraction of non-gapped consensus regions as trusted contigs for elongation with SPAdes (Prjibelski et al., 2020) and II) replacement of gapped regions with the corresponding reference sequence.

Consensus sequences were extracted and polished as above. All the consensus sequences were aligned using MAFFT v7.526 with the *auto* command and then manually inspected in Geneious Prime 2026.0.1 (Biomatters INC., Boston, MA, USA) to obtain the final draft genome.

The assembled genomes were annotated by transferring annotations from appropriate reference sequences using Geneious Prime. For coronaviruses, ORF1a and ORF1ab were annotated manually by identifying the ribosomal frameshift (slippery) sequence TTTAAAC located immediately upstream of the ORF1a stop codon. To generate the final genome sequences, all annotated coding regions were manually inspected to identify and correct erroneous frameshifts introduced during assembly.

### Gap-closing by Sanger sequencing

For some samples, PCRs spanning gaps were produced using primer pairs which were designed based on the draft genomes generated above (**Table 1**). The products were amplified using AccuPrime high-fidelity DNA polymerase (Thermo Scientific, Thermo Fisher Scientific, Waltham, MA, USA) in a final volume of 50 μl using 94°C for 30 s followed by 35 cycles of 94°C for 15 s, 55°C for 30 s and 68°C for 2 min, with a final extension of 68°C for 2 min. The initial PCR products were generated using the respective outer primers, and then used as templates for the nested PCRs with the respective inner primers. The PCR products were purified using the GeneJET PCR Purification Kit (Thermo Scientific) according to the manufacturer’s instructions. The purified products were sequenced using the Sanger system with a combination of BigDye Terminator v. 3.1. Cycle Sequencing Kit (Applied BioSystems, Waltham, MA, USA) with 10 µM primers, purification using Sephadex (Cytiva, Marlborough, MA, USA) and an ABI3500 Genetic Analyzer (Applied BioSystems). Sequences were analyzed using Geneious Prime.

### Phylogenetic analyses

Complete genome sequences belonging to the family *Coronaviridae* (taxID 11118) were downloaded from NCBI (accessed January 22, 2026). To retain only members of *Orthocoronavirinae* and unassigned coronaviruses, sequences classified within the subfamilies *Pitovirinae* (taxID 2946643) and *Letovirinae* (taxID 2501930) were excluded. Due to the substantial overrepresentation of SARS-CoV-2 in public databases, sequences classified as SARS-CoV-2 (taxID 2697049) were also excluded. Instead, SARS-CoV-2 and SARS-CoV-2–related sequences were incorporated from a curated dataset (Tan et al., 2023) and ICTV representative sequences (Woo et al., 2023). The combined dataset was filtered to retain genomes with ≤1% ambiguous nucleotides and with lengths between 21.5 kb and 37 kb. Duplicate sequences were removed. The filtered database was clustered using Vclust v1.3.1 (Zielezinski et al., 2025) at 46% total average nucleotide identity (tANI), followed by subclustering at 85% tANI. Within each subcluster, ICTV reference sequences and previously published sequences (Johnston et al., 2025; Lazov et al., 2021, 2025; Quaade, 2024) were preferentially retained. The total number of sequences per cluster was then limited to approximately 20 genomes, with the remaining sequences randomly selected using Python’s *random.sample* function, ensuring that at least one sequence was retained from each subcluster. This resulted in the CoV-representative dataset.

To place the novel sequences in the context of known CoV diversity, alignment-free pairwise distances based on tANI were computed using Vclust on the CoV-representative dataset together with the novel sequences. Neighbor-joining (Saitou & Nei, 1987) were reconstructed using the *nj* function in the ape package v5.7.1 (Paradis & Schliep, 2019) in R and rooted using delta-CoVs as the outgroup. Genus and subgenus level clades were identified based on the tree topology, and taxonomic annotations were curated accordingly. Where phylogenetic placement was inconsistent with existing annotations, taxonomic assignments were updated to reflect clustering patterns. Based on the curated genus and subgenus level assignments, corresponding clusters were extracted and mapped back to the original dataset. All sequences classified as alpha-CoV were extracted, and a representative dataset was generated as above using Vclust at 80% tANI with subclustering at 91% tANI, limiting each cluster to approximately 25 genomes. This resulted in the alpha-CoV representative dataset.

From the alpha-CoV representative dataset, sequences belonging to *Minacovirus*, *Nyctacovirus*, and *Pedacovirus* were extracted and aligned separately with the relevant novel coronaviruses using MAFFT v7.526 (Katoh & Standley, 2013) with default parameters. Human alpha-CoV 229E (subgenus *Duvinacovirus*) was included as the outgroup to enable subsequent tree rooting. To reduce the impact of poorly aligned and incomplete regions, alignment ends were trimmed to the first and last positions at which the majority (95%) of sequences contained non-Ns. In addition, genome positions where gaps or Ns were assigned to more than 20% of the sequences, were removed from the alignment. Maximum-likelihood trees were inferred using IQ-TREE v2.3.6 (Nguyen et al., 2015) with substitution models determined by ModelFinder (Kalyaanamoorthy et al., 2017). Branch support was assessed using ultrafast boot-strapping (UFBoot) (Hoang et al., 2018) and approximate likelihood-ratio tests (SH-aLRT) (Guindon et al., 2010), each with 1,000 replicates. All trees were visualized in ggtree v3.6.2 (Yu et al., 2017).

For the *Pedacovirus*, genome regions corresponding to the panCoV-B and panCoV-C amplicons were extracted and aligned with MAFFT with default parameters. The alignment included partial sequences obtained by NGS and Sanger sequencing, as well as previously published Danish partial sequences (Lazov et al., 2018). Maximum-likelihood phylogenetic inference was performed as described above.

## Results

### Screening for CoV RNA

Samples from a total of 207 mustelids were collected nationwide (**Figure 1** and **Table 2**). Of the 206 samples screened for SARS-CoV-2, none tested positive. Among the 203 samples screened using pan-CoV assays, one sample from a European polecat (*Mustela putorius*) tested positive, indicating the presence of a coronavirus other than SARS-CoV-2, likely an alphacoronavirus, and was subsequently sequenced by NGS (**Table S1**).

**Table 2:** Sampling overview.

| Order/Family | Species | Common name | Total | Pan-CoV<sup>α</sup> | SARS-CoV-2<sup>α</sup> |
| --- | --- | --- | --- | --- | --- |
| Carnivora/Mustelidae | *Lutra lutra* | Eurasian otter | 60 | 0/60<sup>β</sup> | 0/60<sup>γ</sup> |
| Carnivora/Mustelidae | *Martes foina* | Beech marten | 47 | 0/45<sup>β</sup> | 0/47<sup>γ</sup> |
| Carnivora/Mustelidae | *Martes martes* | European pine marten | 22 | 0/21<sup>β</sup> | 0/21<sup>γ</sup> |
| Carnivora/Mustelidae | *Meles meles* | Euroasian badger | 27 | 0/27<sup>β</sup> | 0/27<sup>γ</sup> |
| Carnivora/Mustelidae | *Mustela erminea* | Stoat | 3 | 0/3<sup>β</sup> | 0/3<sup>γ</sup> |
| Carnivora/Mustelidae | *Mustela nivalis* | Least weasel | 18 | 0/18<sup>β</sup> | 0/18<sup>γ</sup> |
| Carnivora/Mustelidae | *Mustela putorius* | European polecat | 30 | 1/29<sup>β</sup> | 0/30<sup>γ</sup> |
| Chiroptera/Vespertilionidae | *Eptesicus serotinus* | Serotine bat | 11 | 2/11<sup>δ</sup> | 0/11<sup>δ</sup> |
| Chiroptera/Vespertilionidae | *Myotis bechsteinii* | Bechstein's bat | 8 | 0/8<sup>δ</sup> | 0/5<sup>δ</sup> |
| Chiroptera/Vespertilionidae | *Myotis brandtii* | Brandt's bat | 1 | 0/1<sup>δ</sup> | 0/1<sup>δ</sup> |
| Chiroptera/Vespertilionidae | *Myotis dasycneme* | Pond bat | 13 | 4/13<sup>δ</sup> | 0/13<sup>δ</sup> |
| Chiroptera/Vespertilionidae | *Myotis daubentonii* | Daubenton's bat | 71 | 16/70<sup>δ</sup> | 0/71<sup>δ</sup> |
| Chiroptera/Vespertilionidae | *Myotis mystacinus* | Whiskered bat | 2 | 0/2<sup>β</sup> | 0/2<sup>β</sup> |
| Chiroptera/Vespertilionidae | *Myotis nattereri* | Natterer's bat | 7 | 1/7<sup>δ</sup> | 0/7<sup>δ</sup> |
| Chiroptera/Vespertilionidae | *Nyctalus noctula* | Common noctule | 1 | 0/1<sup>β</sup> | 0/1<sup>β</sup> |
| Chiroptera/Vespertilionidae | *Pipistrellus nathusii* | Nathusius' pipistrelle | 2 | 0/2<sup>δ</sup> | 0/2<sup>δ</sup> |
| Chiroptera/Vespertilionidae | *Pipistrellus pipistrellus* | Common pipistrelle | 22 | 5/22<sup>δ</sup> | 0/22<sup>δ</sup> |
| Chiroptera/Vespertilionidae | *Pipistrellus pygmaeus* | Soprano pipistrelle | 57 | 14/57<sup>δ</sup> | 0/56<sup>δ</sup> |
| Chiroptera/Vespertilionidae | *Plecotus auritus* | Brown long-eared bat | 16 | 1/16<sup>δ</sup> | 0/12<sup>δ</sup> |
| Chiroptera/Vespertilionidae | *Vespertilio murinus* | Parti-coloured bat | 7 | 2/7<sup>δ</sup> | 0/7<sup>δ</sup> |
| Chiroptera/ | Not determined |  | 5 | 0/5<sup>δ</sup> | 0/4<sup>δ</sup> |

<sup>α</sup> Samples reactive in screening assay / total screened. <sup>β</sup> Feces. <sup>γ</sup> Throat swabs. <sup>δ</sup> Colon, colon contents, colon swab or fecal samples.

In total, 223 bat samples were obtained through the national surveillance program between 2021 and 2024 (**Figure 2** and **Table 2**). Of these, 214 samples were screened for SARS-CoV-2, and none were positive. Of the 222 samples screened using pan-CoV assays, 45 yielded a positive signal. Positive signals were detected in samples from *Eptesicus serotinus* (*n* = 2), *Myotis dasycneme* (*n* = 4), *Myotis daubentonii* (*n* = 16), *Myotis nattereri* (*n*=1), *Pipistrellus pipistrellus* (*n* = 5), *Pipistrellus pygmaeus* (*n* = 14), *Plecotus auritus (n* = 1), and *Vespertilio murinus* (*n* = 2), as well as six samples identified only as Chiroptera, which was not further typed. Pan-CoV positive samples were subsequently subjected to Sanger sequencing and/or mNGS (**Table S2**). Thirty-five of the pan-CoV–positive bat samples were analyzed by Sanger sequencing. Of these, 13 yielded alpha-CoV sequences based on BLASTn analysis, whereas the remaining samples produced either no matches or no matches of viral origin. Positive Sanger results were obtained from one *P. pygmaeus*, one *M. dasycneme*, and ten *M. daubentonii* samples. None of the samples from *E. serotinus*, *P. auritus*, or *V. murinus* yielded CoV sequences by Sanger.

### Genome recovery and sequence analysis

The pan-CoV positive *M. putorius* sample was subjected to NGS, resulting in the recovery of a full-length alpha-CoV genome with a mean sequencing depth of >150x (**Table 3**).

**Table 3:** Full-length genomes sequences.

| Name | Virus | Length (nt) | Mapped reads | Mean depth<sup>a</sup> | Breadth $\geq 10\times$<sup>b</sup> | Ambiguous bases<sup>c</sup> | Total Ns<sup>d</sup> |
| --- | --- | --- | --- | --- | --- | --- | --- |
| BtCoV/22-8070-56/M.das/DK/2022 | Alphacoronavirus | 27,788 | 5,069 | 17.02 | 80.74% | 2 | 0 |
| BtCoV/22-6959-24/M.dau/DK/2022<sup>e</sup> | Alphacoronavirus | 28,096 | 3,399 | 9.85 | 42.58% | 98 | 347<sup>f</sup> |
| BtCoV/22-8065-78/M.dau/DK/2022<sup>e</sup> | Alphacoronavirus | 28,085 | 78,262 | 183.56 | 99.80% | 0 | 0 |
| BtCoV/22-8071-51/M.dau/DK/2022<sup>e</sup> | Alphacoronavirus | 28,171 | 50,450 | 109.76 | 98.30% | 0 | 58<sup>g</sup> |
| BtCoV/22-8075-42/M.dau/DK/2022 | Alphacoronavirus | 28,217 | 81,639 | 324.85 | 99.95% | 0 | 0 |
| BtCoV/22-8079-53/M.dau/DK/2022<sup>e</sup> | Alphacoronavirus | 28,133 | 8,092 | 21.75 | 78.63% | 171 | 175<sup>h</sup> |
| BtCoV/22-8084-55/M.dau/DK/2022 | Alphacoronavirus | 28,129 | 13,136 | 51.13 | 99.50% | 0 | 0 |
| BtCoV/OV2024-44/P.pyg/DK/2024<sup>e</sup> | Alphacoronavirus | 27,918 | 87,213 | 303.88 | 100.00% | 1 | 0 |
| FRCoV/24-10470-79/M.put/DK/2024 | Alphacoronavirus | 28,508 | 51,042 | 192.49 | 100.00% | 6 | 0 |

<sup>a</sup> Mean sequencing depth across the genome. <sup>b</sup> Percentage of genome positions covered by $\geq 10\times$ sequencing depth. <sup>c</sup> Number of ambiguous nucleotides, excl. N. <sup>d</sup> Total number of undetermined nucleotides (N) in the final consensus sequence, including assembly gaps. <sup>e</sup> Deep sequenced (see methods). <sup>f</sup> Eleven putative gaps ranging from 4-91 nt located across the genome, incl. Spike. <sup>g</sup> Two gaps for 23 nt and 35 nt located in ORF1a. <sup>h</sup> Five gaps ranging from 3-88 nt, located in ORF1ab and Spike.

In total, 53 samples from bats were also subjected to NGS, including a subset of pan-CoV negative samples, in order to explore the presence of other viral sequences (**Table S2**). Six full-length alpha-CoV genomes were obtained from *M. daubentonii*, one from *M. dasycneme*, one from *P. pygmaeus* (feces from several individuals) (**Table 3**, **Figure S1**). An additional six *M. daubentonii* samples yielded fewer than 1000 alpha-CoV reads, precluding genome assembly (**Table S3**). However, in two of these samples (22-8066-62 and 22-8071-52), partial sequences corresponding to the panCoV-B and/or panCoV-C amplicons in ORF1b were recovered, despite the low read counts.

All Sanger-positive samples contained more than 600 alpha-CoV reads by NGS, with the exception of the *P. pygmaeus* sample 23177-28, which produced only four alpha-CoV reads. Conversely, the *M. daubentonii* samples 22-8083-63 and 22-8068-70 did not yield alpha-CoV sequences by Sanger, but produced 339 and 709 alpha-CoV reads by NGS, respectively. Sanger sequencing was not performed on the *P. pygmaeus* sample OV2024-44, which tested positive in the pan-CoV assay; however, NGS resulted in recovery of a full-length alpha-CoV genome.

One *M. dasycneme*, the single *M. nattereri*, one *P. pipistrellus* and six *P. pygmaeus* samples that were positive in the pan-CoV assay did not yield alpha-CoV reads by NGS. However, the *P. pipistrellus* sample (22-7401) and two of the *P. pygmaeus* samples (2022-7606 and 2024-00582-42) contained other viral sequences, including astrovirus, betapaprhavirus, picorna-like virus, and permutotetra-like virus. Seven full-length astrovirus genomes were recovered from the bats: Six from *M. daubentonii* and one from the *P. pipistrellus* mentioned above (**Table S4**). In samples where astrovirus was detected, co-infection with alpha-CoV was observed, except for the *P. pipistrellus* sample (22-7401).

Of the 16 samples subjected to NGS despite being negative in the pan-CoV assay, seven contained viral sequences, including unclassified *Riboviria* species, *iflavirus*, and *permutotetra*-like viruses. The latter two are insect-associated viruses and were likely present due to dietary origin.

In summary, alpha-CoVs were detected across multiple host species. In *M. daubentonii*, twelve samples were found positive for alpha-CoVs by Sanger and/or NGS, including six full-length genomes. In *P. pygmaeus*, two samples were positive, including one yielding a full-length genome. In addition, full-length alpha-CoV genomes were recovered from one *M. dasycneme* and one *M. putorius*.

### Phylogenetic analyses

A global phylogenetic tree based on alignment-free pairwise distances revealed that the sequence obtained from *M. putorius* belonged to the subgenus *Minacoviru*s, whereas the seven genomes obtained from *M. daubentonii* and *M. dasycneme* belonged to the subgenus *Pedacovirus* (**Figure 3**). The genome obtained from *P. pygmaeus* clustered within the subgenus *Nyctacovirus*. The phylogenetic analysis further identified several discrepancies between existing NCBI annotations and tree topology (**Table S5**). Local alignment and rooted maximum-likelihood phylogenetic inference were subsequently performed separately for each of the three subgenera (**Table S6**).

**Figure 3.**
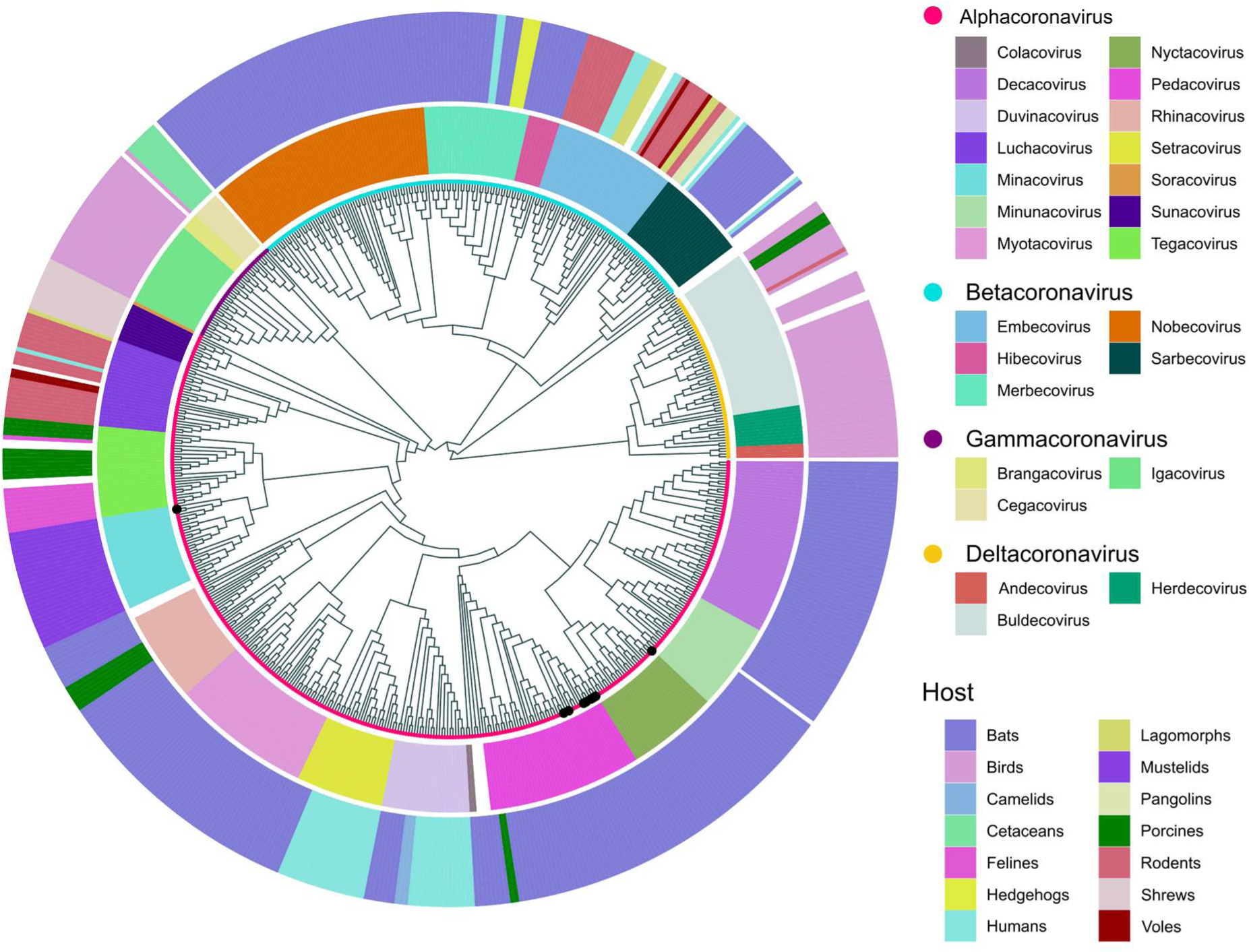
Alignment-free phylogeny of global coronavirus genome diversity (n = 628), including the novel coronavirus genomes. The inner ring denotes viral genus (novel genomes denoted with black circles), while the middle and outer rings denote viral subgenus and broader host groups, respectively.

Within *Minacovirus*, the novel *M. putorius* viral genome formed a distinct clade together with other viruses isolated from *M. putorius* and M. *putorius furo* (domesticated from *M. putorius*), clearly separated from the viral genomes identified in *N. vison* (**Figure 4**). Within this clade, the novel genome clustered most closely together with a *M. putorius* sequence isolated in the Netherlands in 2010, sharing 90.6% nucleotide sequence identity.

**Figure 4.**
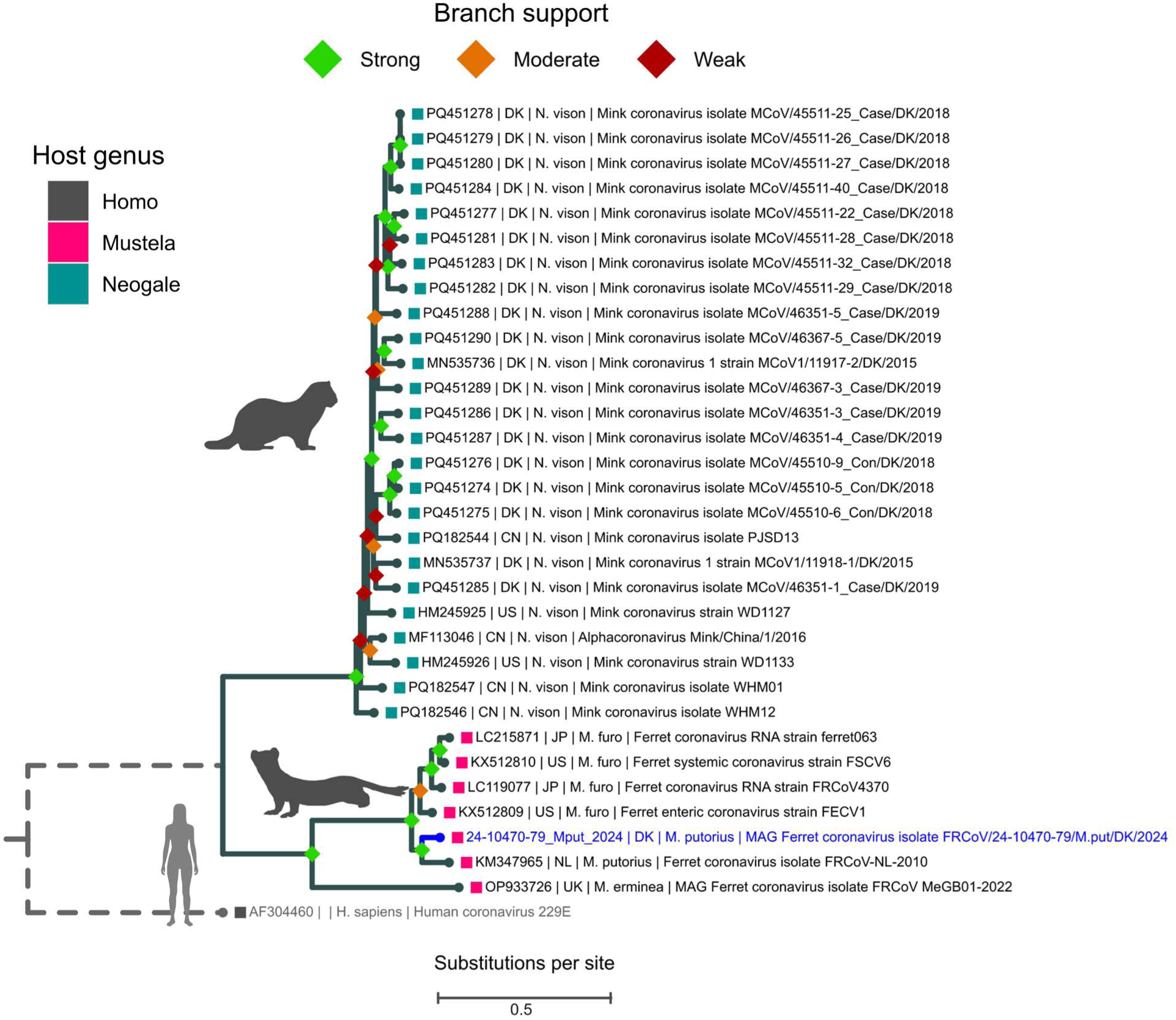
Rooted maximum-likelihood tree of full-length nucleotide sequences of minacoviruses with the novel virus obtained from a Danish *Mustela putorius* in blue and the outgroup in grey. Branch support is categorized as strong (SH-aLRT ≥ 80 and UFBoot ≥ 95), moderate (SH-aLRT ≥ 60 and UFBoot ≥ 60), and weak (below these thresholds). Phylogenetic inference was performed in IQ-TREE using the GTR+F+I+G4 substitution model with 1,000 replicates for both SH-aLRT and UFBoot. Solid branches are proportional to the scale bar (substitutions per site), while dashed branches have been truncated for visualization purposes.

In the *Nyctacovirus* phylogeny, the novel *P. pygmaeus* viral genome grouped together with previously reported *P. pygmaeus* viral genomes from Denmark and Sweden (**Figure 5**), sharing 96.4-98% nucleotide sequence identity. This clade was nested within a larger clade that also included a viral genome obtained from *Pipistrellus kuhlii* in Italy, and one from *Afronycteris nana* in Eswatini. Comparison with the previously reported Danish strain (accession no. MN482242) revealed a total of 54 amino acid (aa) differences across the genome. The majority of these non-synonymous substitutions were located in ORF1a (19 aa) and Spike (29 aa). In contrast, ORF1b, ORF3 and N each contained a single aa substitution, while M contained three aa differences, including one insertion immediately downstream of the start codon. No amino acid differences were observed in the E protein.

**Figure 5.**
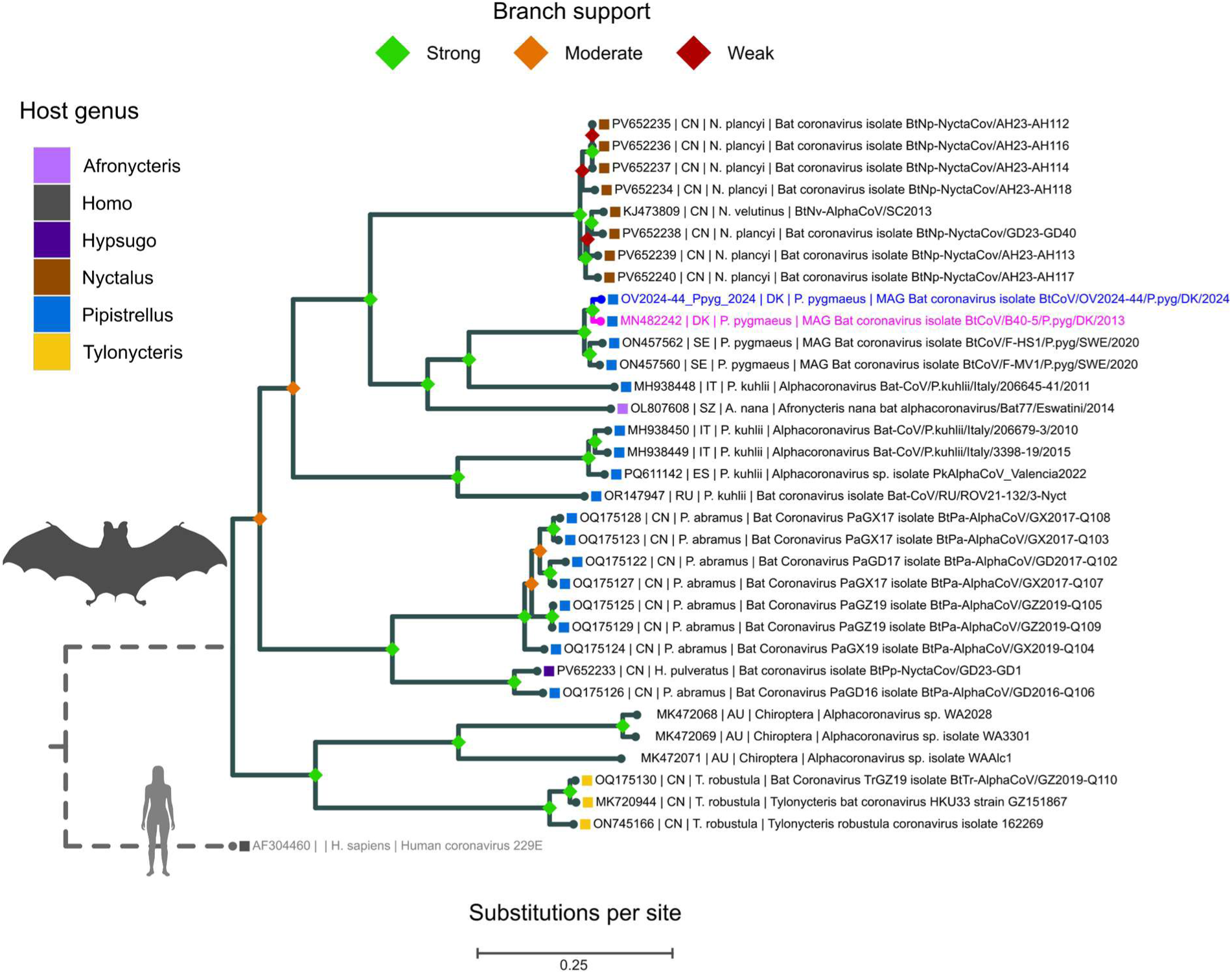
Rooted maximum-likelihood tree of full-length nucleotide sequences of nyctacoviruses, with the novel virus obtained from a Danish *Pipistrellus pygmaeus* shown in blue, previously reported Danish sequences in magenta, and the outgroup in grey. Branch support is categorized as strong (SH-aLRT ≥ 80 and UFBoot ≥ 95), moderate (SH-aLRT ≥ 60 and UFBoot ≥ 60), and weak (below these thresholds). Phylogenetic inference was performed in IQ-TREE using the GTR+F+I+G4 substitution model with 1,000 replicates for both SH-aLRT and UFBoot. Solid branches are proportional to the scale bar (substitutions per site), while dashed branches have been truncated for visualization purposes.

For *Pedacovirus*, all seven novel viral genomes from *M. daubentonii* and *M. dasycneme* clustered within the same major clade (**Figure 6**). Five of the *M. daubentonii* viral genomes formed a well-supported subclade together with previously reported *M. daubentonii* viral genomes from Denmark, Finland, United Kingdom and Spain, sharing 93.7-96.9% sequence identity. The remaining *M. daubentonii* viral genomes grouped within a sister clade containing four other *M. daubentonii* viral genomes, three from Denmark and one from United Kingdom, sharing 93.4-93.9% nucleotide sequence identity. In contrast, the *M. dasycneme* viral genome clustered together with sequences obtained from *Pipistrellus* species from Denmark, United Kingdom, the Netherlands, Russia and Sweden, and a *Hypsugo savii* viral genome from Italy, sharing only 83.4-88.2% sequence identity.

**Figure 6.**
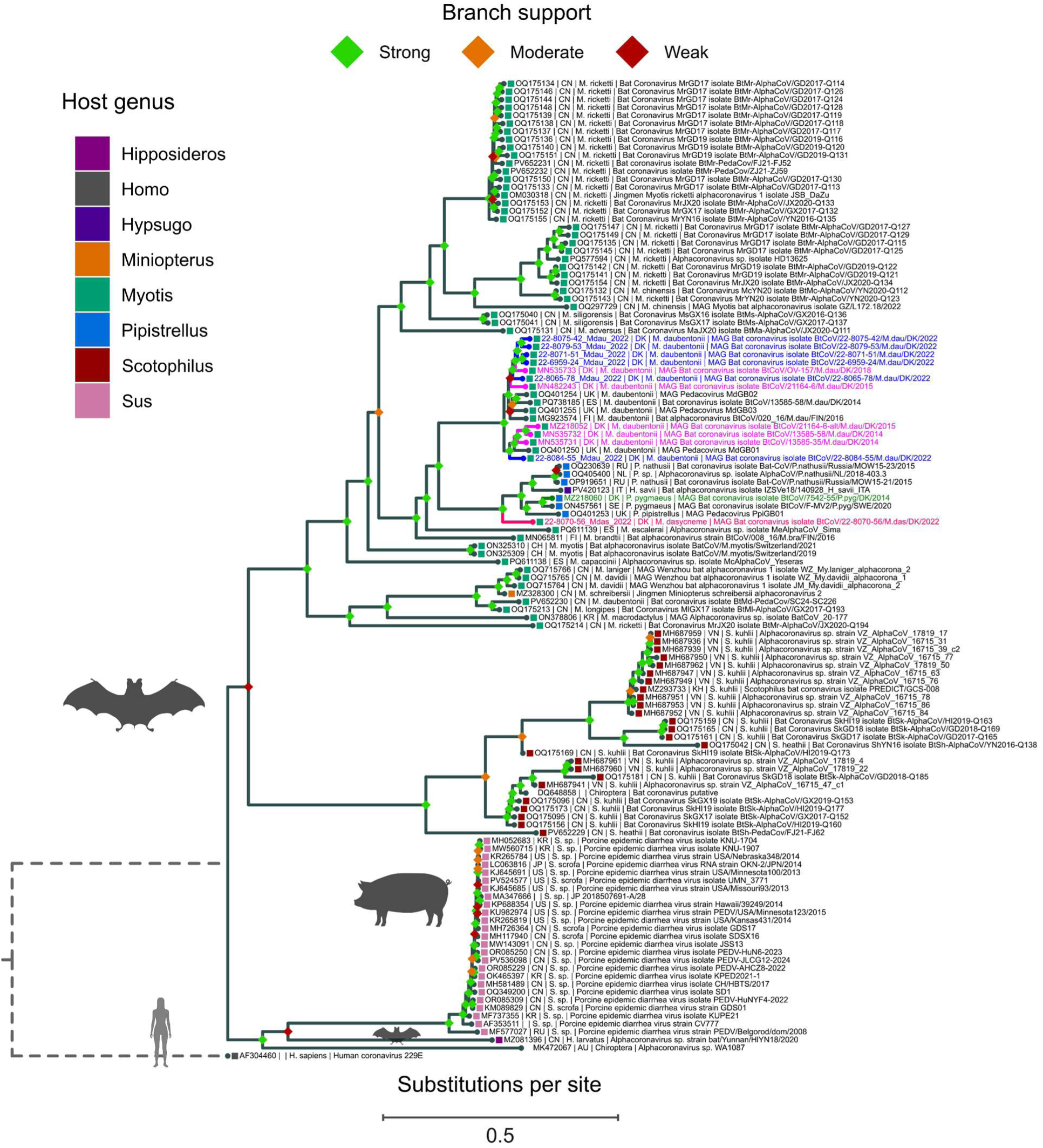
Rooted maximum-likelihood tree of full-length nucleotide sequences of pedacoviruses, with novel viruses obtained from Danish *Myotis daubentonii* and *Myotis dasycneme* shown in blue and hot pink, respectively; previously reported Danish *M. daubentonii* sequences in magenta; a previously reported *Pipistrellus pygma*eus sequence in green; and the outgroup in grey. Branch support is categorized as strong (SH-aLRT ≥ 80 and UFBoot ≥ 95), moderate (SH-aLRT ≥ 60 and UFBoot ≥ 60), and weak (below these thresholds). Phylogenetic inference was performed in IQ-TREE using the GTR+F+I+G4 substitution model with 1,000 replicates for both SH-aLRT and UFBoot. Solid branches are proportional to the scale bar (substitutions per site), while dashed branches have been truncated for visualization purposes.

To place the newly recovered genomes into the context of previous Danish findings, where only short panCoV-B and panCoV-C fragments were available, additional maximum-likelihood trees were inferred for these regions (130 bp and 208 bp, respectively). These analyses included partial viral sequences from the *M. daubentonii* samples 22-8066-62 (panCoV-B) and 22-8071-52 (panCoV-B and C), together with previously published viral sequences and reference datasets (**Table S7**, **Table S8**). Consistent with whole genome phylogeny, both samples (obtained from *M. daubentonii*) clustered with previously reported *M. daubentonii* viral sequences, as well as a single *M. dasycneme* viral sequence from Denmark (**Figure S2**, **Figure S3**). However, due to the short alignment lengths, branch support values were generally weak, and the resulting trees should be interpreted with caution.

## Discussion

In this study, bat and mustelid samples collected through national surveillance were screened for alpha- and beta-CoVs, including SARS-CoV-2, and positive samples were further processed using an optimized metagenomic workflow to enable recovery and characterization of full-length viral genomes. None of the 214 bat samples or 206 mustelid samples tested positive for SARS-CoV-2. Following PCR screening and subsequent confirmation by Sanger sequencing and/or mNGS, alpha-CoVs were detected in 15 of the 222 bat samples and in one of the 203 mustelid samples analyzed. Among the bat samples, one positive originated from *M. dasycneme,* twelve from *M. daubentonii*, and two from *P. pygmaeus*. Full-length genomes were recovered from the *M. dasycneme* sample, six *M. daubentonii* samples, and one *P. pygmaeus* sample, as well as partial genomes from two additional *M daubentonii* samples. In addition, a full-length genome was obtained from the alpha-CoV positive mustelid sample originating from *M. putorius*.

The detection of alpha-CoVs in bats in this study is consistent with previous findings from Denmark (Lazov et al. 2018 and 2021), and further supports the endemic presence of these viruses in Danish bat populations. In contrast, no beta-CoVs were detected in the bats during the study period, despite the prior identification of a MERS-related betacoronavirus in *P. auritus* in Denmark in 2020 (Johnston et al., 2025). This discrepancy may reflect temporal variation in virus circulation, low prevalence, limited sample size, or differences in sampling strategy.

Seventeen species of bats have been registered in Denmark (Baagøe, 2007) all belonging to the family Vespertilionidae. The limestones mines of Daugbjerg and Mønsted, which are located only a few kilometers apart in western Jutland, are important hibernacula for *Myotis* bats. Each autumn, thousands of bats from large parts of Jutland gather at these sites to spend the winter in hibernation (Baagøe, 2001; Baagøe & Degn, 2009; Elmeros et al., 2022) (Baagøe, 2001; Baagøe & Degn, 2009; Elmeros et al., 2022). *M. daubentonii* is the most numerous species (19.000), followed by some 6.000 *M. dasycneme* and no more than 100-200 *M. brandtii* and *M. natteri*. The full-length viral genomes obtained in this study from *M. daubentonii* as well as those reported previously by Lazov et al. 2021 originate from bats sampled at Mønsted or Daugbjerg, and cluster within the same *Pedacovirus* clade. A similar pattern was observed for the partial ORF1b sequences corresponding to the panCoV-B and panCoV-C regions. This clade appears to be almost exclusively composed of sequences from *M. daubentonii*, with only a single partial sequence derived from *M. dasycneme*. This pattern suggests that the circulating pedacoviruses in this region, may be strongly associated with *M. daubentonii*, potentially reflecting host-specific transmission within the large underground hibernacula.

The *M. dasycneme* genome, also obtained from Mønsted, belonged to a separate *Pedacovirus* clade, clustering with sequences obtained from *Pipistrellus* species from Denmark, the United Kingdom, the Netherlands, Russia, and Sweden, and a *Hypsugo savii* from Italy, sharing only 83.4–88.2% nucleotide sequence identity. The Danish *P. pygmaeus* sequence in this lineage was previously isolated in 2014 in Vadum, located approximately 85 km northeast of the Mønsted limestone mines. Unlike the *Myotis* bats, *P. pygmaeus* has never been found in underground hibernacula in northern Europe. Occasional close contact to other bat species is likely to occur in tree holes and buildings, which are the preferred roost sites for *P. pipistrellus* all year round. Compared with the largely homogeneous and host-associated clades observed for *M. daubentonii*, this lineage appears considerably more heterogeneous, both genetically and with respect to host species. This pattern may indicate that certain *Pedacovirus* lineages circulate across multiple vespertilionid hosts. The apparent lack of host structure could reflect limited sampling of this lineage, as relatively few genomes have been obtained compared with those from *M. daubentonii*. Additional sampling across host species and geographic regions will therefore be required to determine whether this clade represents a broader host range or simply an under-sampled lineage.

Some bat species in Europe, such as *Pipistrellus nathusii, Nyctalus noctula*, and *Vespertilio murinus*, are known to migrate long distances in autumn and spring. Others, e.g. *M. nattereri, Plecotus auritus, and E. serotinus,* are rated as much more sedentary, with movements between summer and winter habitats of generally max. 50–100 km. Species like *M. daubentonii* and *M. dasycneme* are rated as facultative migrants covering intermediate migrating distances up to 300–400 km (Baagøe, 2001; Encarnação & Becker, 2023; Gaisler et al., 2003; Haarsma, 2023; Hutterer et al., 2005) probably depending on distances between suitable summer habitats, and known hibernation and swarming sites. More recent research indicates that there is a substantial variation in migration distances within species.

Most of this information is based on recovery of relatively few ringed individuals, and the exact summer or winter habitats are not always known, so migration distances might be significantly longer. Of the several hundred bats of four species ringed in the Danish limestone mines in the 1950’s and 60’s (Egsbæk & Jensen, 1963) none were recaptured in northern Germany and likewise, bats ringed in northern Germany have never been found in the Danish limestone mines. However, during the intensive population research in Mønsted in 2009 (Baagøe & Degn, 2009) a single *M. dasycneme* female was found that had been ringed as a juvenile the year before near Kiel in northern Germany (F. Gloza-Rausch, pers. comm.). The following year, this individual was hibernating in northern Germany near Itzehoe, and the following summer, it was back in the colony where it was born (P. Borkenhagen pers. com.).

Furthermore, it has been shown that in late summer and autumn, both *M. daubentonii* and *M. dasycneme* can be frequently observed both hunting insects and in more directional flight far at sea in the Baltic (Ahlén et al., 2009). *M. daubentonii* is one of the species that each year in late summer and autumn gathers at certain points on the coasts of southern Sweden and southern Denmark (Lolland-Falster) and can be seen flying out from these points directly over the sea on what seems to be regular migration. Such *M. daubentonii* individuals in unidirectional flight have been observed in the middle of the Baltic Sea and also migrating out from the south-western point of Bornholm (Ahlén et al., 2009; Baagøe, 2011; Møller et al., 2013).

*M. daubentonii* is a very common bat distributed widely and continuously over large parts of the Nordic countries, the Baltic states, Poland and Germany (Baagøe, 2007; Borkenhagen, 2011; Encarnação & Becker, 2023) presumably with breeding colonies, intermediate day roosts, etc. all over the region. In the active part of the year, many bats often roost together, in confined spaces. In such situations they are often fully active, creating favorable conditions for fecal-oral virus transmission, and frequent contact between individuals from neighboring roosting sites is likely.

There is evidence for substantial gene flow between colonies across western continental Europe for *M. daubentonii* (Encarnação & Becker, 2023). The isolation by distance is stronger for females than males suggesting a wider dispersal for males. Microsatellite markers suggest gene flow between pond bats hibernating in Denmark and northern Germany, and pond bat from Danish hibernacula and northern Germany cluster together compared to populations further east and south (Andersen et al., 2019).

In summary, there is migration and more flux between local *M. daubentonii* (and *M. dasycneme*) populations within their ranges in Northern Europe, but we do not know the extent of such flux.

For *M. daubentonii* migration and dispersal may, at least partly, explain why viruses from Denmark and Finland are closely related. However long-term viral persistence in local, stable host populations is likely to play a major role. Most certainly the latter must be the case concerning the similar sequences from the UK, since at least up till now there have been no records of dispersal of *M. daubentonii* between continental Europe and the British Isles.

The *Nyctacovirus* viral genome recovered from *P. pygmaeus* in this study clustered closely with previously reported genomes from Denmark and Sweden, sharing 96.4–98% nucleotide sequence identity. The earlier Danish genome was obtained in 2013 from Sollerup on the island of Fyn, while the genome reported here was sampled in 2024 from Svendborg, located less than 25 km away. Despite the eleven-year interval, the two Danish genomes differed by only 54 amino acids across the genome. The majority of these substitutions were located in ORF1a and Spike, while the remaining genes were highly conserved. This high level of similarity suggests that a relatively stable *Nyctacovirus* lineage has been circulating in Danish *P. pygmaeus* populations for more than a decade. The close relationship to sequences detected in Sweden further indicates that this lineage may be widely distributed among northern European *P. pygmaeus* populations. Together, these observations are consistent with a host-associated alphacoronavirus that persists within *P. pygmaeus* populations over extended periods, potentially maintained through long-term circulation within regional bat populations rather than frequent cross-species transmission.

Summer and winter roosts of *P. pygmaeus* are found in buildings or tree holes (Baagøe, 2007; Jones & Froidevaux, 2023). Individuals may roost singly or in clusters sometimes many individuals together. The breeding colonies in summer often consist of many females and their young in close clusters in well-insulated places often with confined space and ideal conditions for fecal-oral transmission of viruses. In the winter months *P. pygmaeus* hibernate singly or in clusters in cool but frost-free places in protected sites in buildings or tree holes (Baagøe, 2007; Elmeros et al., 2024; Møller et al., 2013). Summer and winter habitats are often located in the same local area. However, *P. pygmaeus* also shows migratory traits, and low levels of genetic structuring across much of the European range suggest large-scale movements (Jones & Froidevaux, 2023). Movements exceeding 100 km, and up to 700 km, have been documented, and genetic studies suggest that females are more likely than males to remain in or return to their natal area (Dietz et al., 2009; Jones & Froidevaux, 2023).

*P. pygmaeus* is the most common bat species in Denmark and southern Sweden occurring almost everywhere (Baagøe, 2007; De Jong et al., 2020; Elmeros et al., 2024). *P. pygmeaus* is frequently observed foraging and migrating over the sea especially in the Baltic area (Ahlén et al., 2009; Smeele et al., 2026) and frequent flux between Danish and southern Swedish populations with ample possibilities for exchange of viruses are highly probable. However, the existence of long-term viral persistence in local, stable host populations cannot be excluded, especially perhaps in marginal parts of the distribution area.

The detection of an alpha-CoV in *M. putorius* represents, to our knowledge, the first such finding in a wild mustelid in Denmark, expanding the known host range of coronaviruses in Danish wildlife and highlighting the importance of continued surveillance across animal populations. Phylogenetically, the genome belongs to the subgenus *Minacovirus* and clustered most closely with a sequence previously detected in a wild European polecat from the Netherlands and with viruses identified in domesticated ferrets (*M. putorius furo*), forming a well-defined clade distinct from the *Minacovirus* lineage circulating in farmed American mink (*N. vison*). This separation was particularly notable given that *Minacovirus* has previously been detected in farmed American mink in Denmark (Lazov et al., 2025; Quaade et al., 2026). The clear phylogenetic distinction between the Mustela-associated lineage and the mink viruses suggests that spillover between farmed mink and wild polecats has not occurred, at least among the sampled animals.

Interestingly, despite the widespread circulation of SARS-CoV-2 in Danish mink farms during the COVID-19 pandemic, none of the wild mustelid samples analyzed in this study tested positive for SARS-CoV-2. Similar findings were reported by Boklund et al., with no detection of SARS-CoV-2 in wildlife sampled near infected farms in Denmark in 2020. Together, these observations may indicate limited transmission from farmed to wild mustelid populations, although continued surveillance is required to fully assess the risk.

The presence of alphacoronaviruses in wild mustelids nonetheless highlights that these animals can act as natural hosts of alpha-CoVs such as *Minacovirus*. Because mustelids occupy ecological niches at the human–animal interface and are known to be susceptible to multiple coronaviruses, their role in coronavirus ecology warrants continued monitoring. Although alpha-CoVs from the subgenus *Minacovirus* are currently not known to pose a direct threat to human health, the occurrence of coronaviruses in non-bat mammalian hosts may be relevant for understanding potential intermediate hosts in future spillover events.

The interpretation of the metagenomic data generated in this study is subject to several methodological considerations related to genome assembly and the broader detection of viral diversity. Accurate genome reconstruction is critical, as errors introduced during assembly can propagate into downstream analyses such as phylogenetic inference and genome annotation (Denton et al., 2014). In metagenomic datasets, de novo assembly typically produces contigs that must subsequently be ordered and connected into longer scaffolds, which may still contain unresolved gaps (Liao et al., 2019). However, uneven sequencing depth, sequencing errors, and compositional biases can produce fragmented assemblies or complex de Bruijn graphs that complicate contig resolution (Liao et al., 2019; Rodrigue et al., 2009).

Reference-guided approaches can help overcome fragmentation by mapping reads or contigs to a related genome. Nevertheless, such approaches introduce the risk of reference bias, where divergent regions may be misaligned, collapsed, or excluded if they differ substantially from the reference sequence (Brandt et al., 2015; Lunter & Goodson, 2011). This limitation can be particularly pronounced when short contigs are mapped to a reference scaffold, as contigs originating from highly divergent regions may fail to align, and therefore be excluded from the reconstructed genome. In coronaviruses, this issue is especially relevant for the Spike gene, which is known to be among the most variable regions of the genome. As a result, some genomic regions may remain difficult to reconstruct accurately despite the use of combined assembly strategies.

Although the primary objective of this study was to characterize coronaviruses, the metagenomic approach also revealed the presence of additional viral sequences in several samples. In particular, full-length astrovirus genomes were recovered from multiple bats, and several samples contained sequences related to insect-associated viruses such as iflaviruses and permutotetra-like viruses. Notably, astrovirus was detected together with alpha-CoV in several samples, indicating that co-infections may be relatively common in these bat populations. Because the focus of the present study was on coronavirus diversity and genome characterization, these additional viral sequences were not analyzed further. Nevertheless, their detection highlights the capacity of metagenomic sequencing to capture broader viral diversity within wildlife samples, even when sampling strategies are primarily designed to target a specific virus group.

## Conclusions

Overall, this study expands our understanding of coronavirus diversity in Danish wildlife. Alphacoronaviruses were detected in several bat species, including *M. daubentonii*, *M. dasycneme*, and *P. pygmaeus*, as well as in a wild mustelid (*M. putorius*). The observed phylogenetic patterns suggest that some CoV lineages, such as the pedacoviruses detected in *M. daubentonii* and the nyctacoviruses in *P. pygmaeus*, may represent relatively stable host-associated lineages circulating within northern European bat populations. In contrast, other lineages appear more heterogeneous and may involve multiple host species, although additional sampling will be required to clarify these patterns. The detection of a *Minacovirus* in the wild European polecat represents the first such finding in Denmark and indicates that alpha-CoVs circulate naturally in wild mustelids independently of those previously identified in farmed mink. Importantly, no evidence of SARS-CoV-2 infection was detected in the surveyed wildlife populations despite extensive circulation in Danish mink farms during the COVID-19 pandemic. Together, these findings emphasize the importance of continued wildlife surveillance and genome-level characterization of coronaviruses to improve our understanding of virus diversity, host associations, and the ecological conditions that may facilitate the emergence of novel coronaviruses.

## Declarations

### Ethical approval

Ethical review and approval were waived for this study. Bat samples consisted of fecal material collected without causing harm to the animals and were obtained under dispensation MST-850-00116 and SN 302-009 from the Danish Environmental Protection Agency. Samples from the passive surveillance of bats and from mustelids were collected exclusively from animals that were already deceased.

### Consent for publication

Not applicable.

### Data availability

MiSeq and NextSeq data are available at NCBI BioProject with accession no. PRJNA1461863. The assembled genomes have been deposited in GenBank under accession numbers XXXX-XXXX. The datasets used and/or analyzed during the current study are available from the corresponding author on reasonable request.

### Authors’ contributions

CMJ and TBR conceived and designed the study. HJB, ME, ETF and MLQ conducted the sampling. Pan-CoV and SARS-CoV-2 screening of mustelids was led by MJSH and TBR. Pan-CoV and SARS-CoV-2 screening of bats was led by TBR. Metagenomics workflow was established by CMJ and VG. Phylogenetic analyses were performed by CMJ. Original draft preparation was performed by CMJ. All authors contributed to reviewing and editing the manuscript. Supervision was performed by LL and TBR. All authors read and approved the final manuscript.

## Acknowledgements

We thank Jani Christiansen, Rasmus Oskar Rask Hansen, and Fie Fisker Brønnum Christensen for their invaluable technical assistance during this study. We also thank Anne Sofie Vedsted Hammer and Tim Kåre Jensen for their contributions to the examinations of bat carcasses. We are grateful to the many volunteers for handing in mustelid carcasses. We further thank Martin Schou Pedersen for assistance with NextSeq sequencing.

## Supplementary

**Table S1**: Mustelid sample collection details.

**Table S2**: Bat sample collection details.

**Table S3:** Partial genome mappings

**Table S4:** Full-length astroviruses

**Table S5:** Coronavirus representative dataset

**Table S6:** *Minacovirus*, *Nyctacovirus* and *Pedacovirus* representative dataset

**Table S7:** Pedaco panCoV-B dataset

**Table S8:** Pedaco panCoV-C dataset

**Figure S1.**
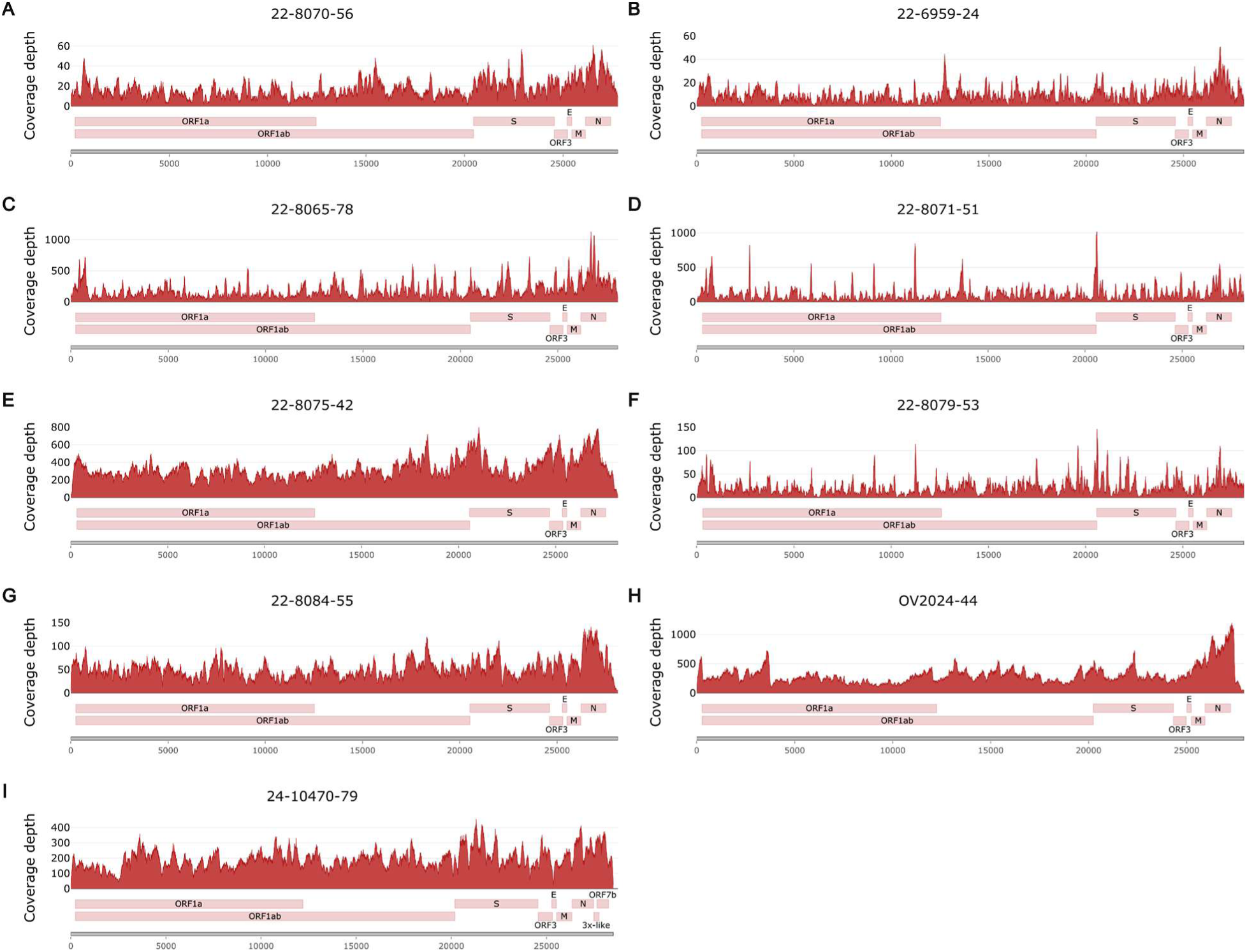
Coverage profiles and genome organization of novel alphacoronaviruses obtained from bats and a mustelid in Denmark. Coverage plots show read depth across the genome, and genome organization diagrams indicate annotated ORFs. A) BtCoV/22-8070-56/M.das/DK/2022, B) BtCoV/22-6959-24/M.dau/DK/2022, C) BtCoV/22-8065-78/M.dau/DK/2022, D) BtCoV/22-8071-51/M.dau/DK/2022, E) BtCoV/22-8075-42/M.dau/DK/2022, F) BtCoV/22-8079-53/M.dau/DK/2022, G) BtCoV/22-8084-55/M.dau/DK/2022, H) BtCoV/OV2024-44/P.pyg/DK/2022, and I) FRCoV/24-10470-79/M.put/DK/2024.

**Figure S2.**
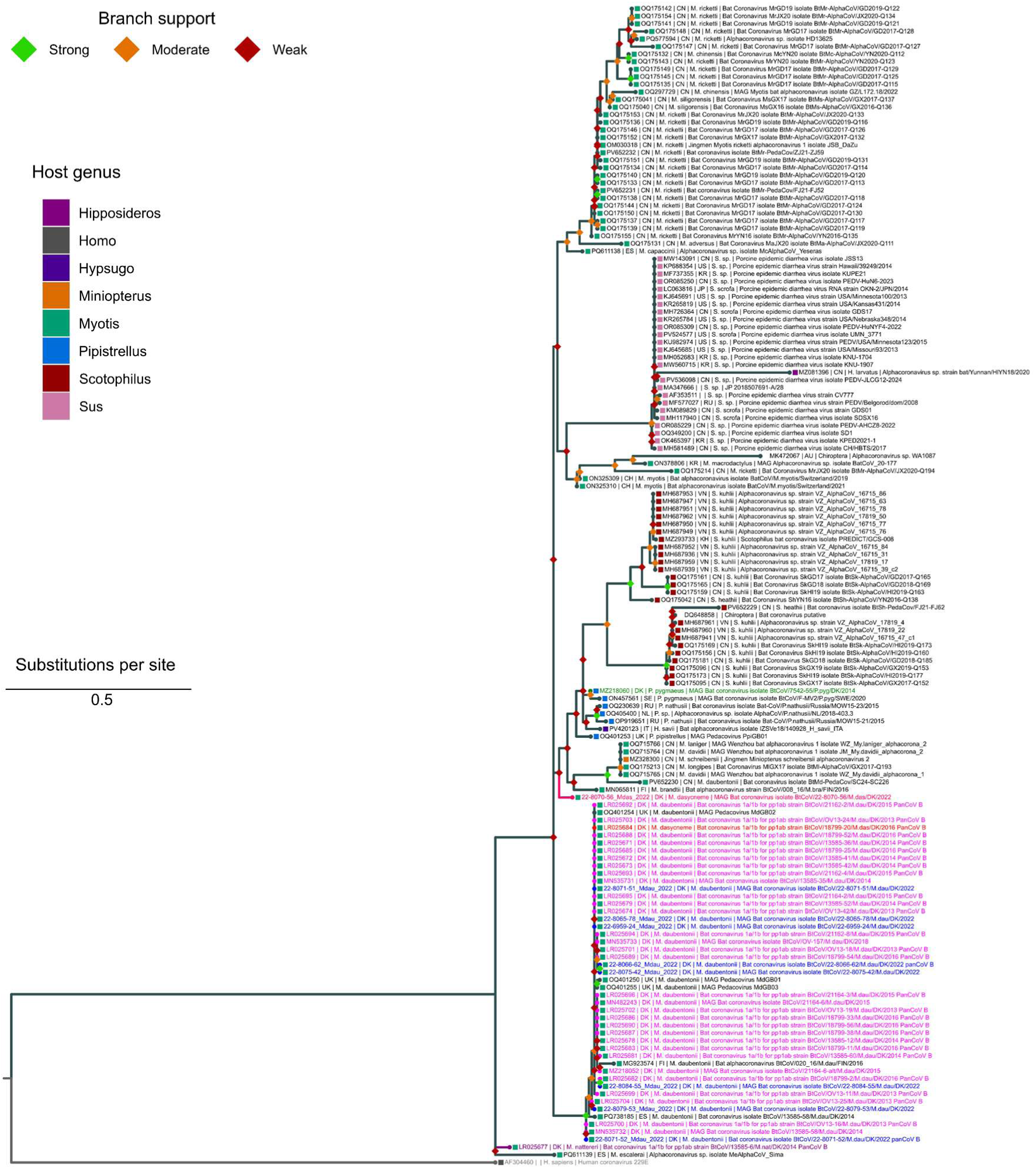
Rooted maximum-likelihood tree of partial nucleotide sequences of pedacoviruses, corresponding to panCoV-B amplicons, with novel viruses obtained from Danish *Myotis daubentonii* and *Myotis dasycneme* shown in blue and hot pink, respectively; previously reported Danish *M. daubentonii* sequences in magenta; a previously reported *M. dasycneme* in red; a previously reported *Pipistrellus pygmaeus* sequence in green; a previously report *Myotis nattereri* in purple; and the outgroup in grey. Branch support is categorized as strong (SH-aLRT ≥ 80 and UFBoot ≥ 95), moderate (SH-aLRT ≥ 60 and UFBoot ≥ 60), and weak (below these thresholds). Phylogenetic inference was performed in IQ-TREE using the HKY+F+G4 substitution model with 1,000 replicates for both SH-aLRT and UFBoot.

**Figure S3.**
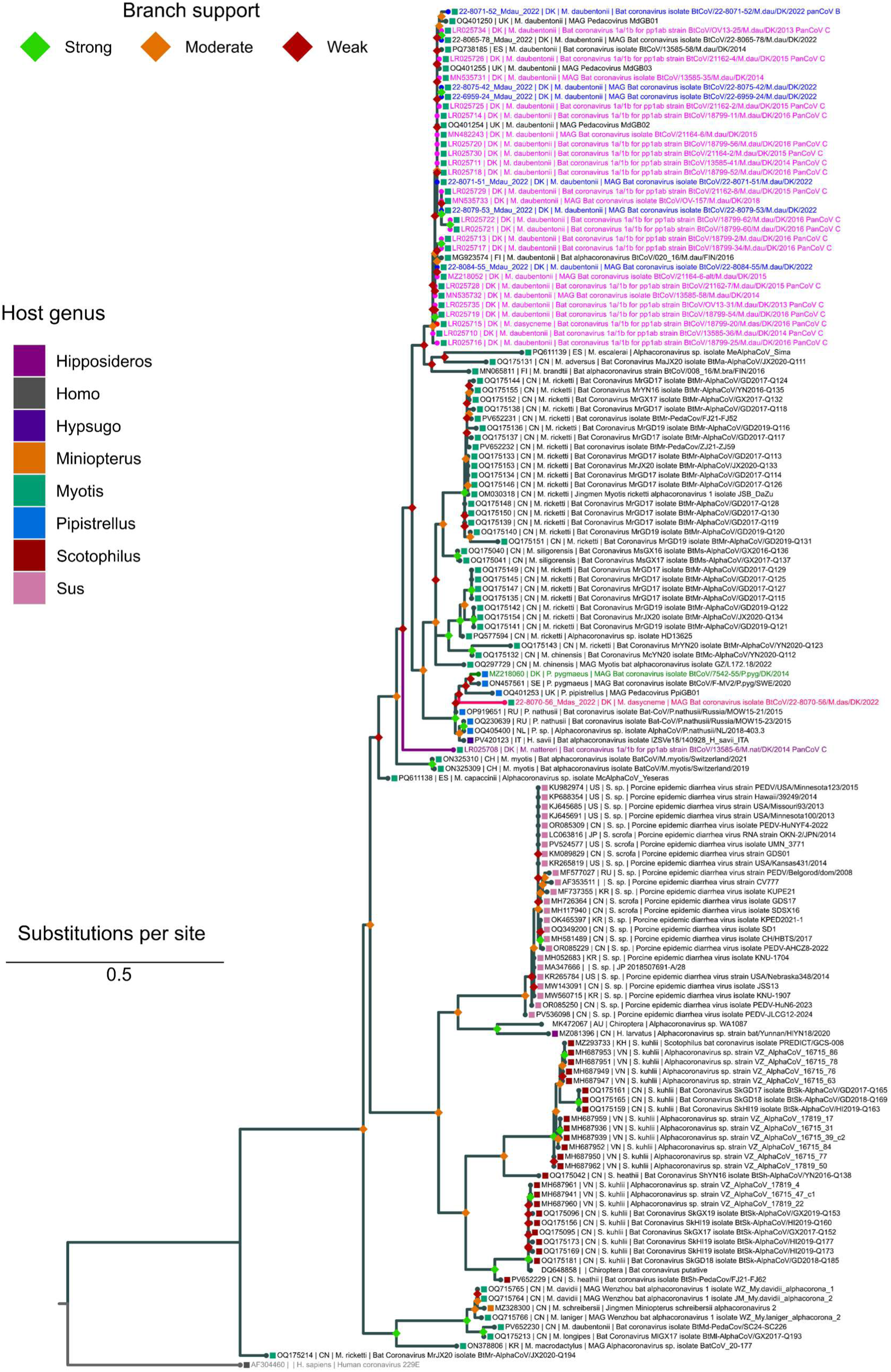
Rooted maximum-likelihood tree of partial nucleotide sequences of pedacoviruses, corresponding to panCoV-C amplicons, with novel viruses obtained from Danish *Myotis daubentonii* and *Myotis dasycneme* shown in blue and hot pink, respectively; previously reported Danish *M. daubentonii* sequences in magenta; a previously reported *M. dasycneme* in red; a previously reported *Pipistrellus pygmaeus* sequence in green; a previously report *Myotis nattereri* in purple; and the outgroup in grey. Branch support is categorized as strong (SH-aLRT ≥ 80 and UFBoot ≥ 95), moderate (SH-aLRT ≥ 60 and UFBoot ≥ 60), and weak (below these thresholds). Phylogenetic inference was performed in IQ-TREE using the HKY+F+G4 substitution model with 1,000 replicates for both SH-aLRT and UFBoot.

